# A Multistaged Hyperparallel Optimization of the Fuzzy-Logic Mechanistic Model of Molecular Regulation

**DOI:** 10.1101/2020.09.28.315986

**Authors:** Paul Aiyetan

## Abstract

**Motivation:** Although it circumvents hyperparameter estimation of ordinary differential equation (ODE) based models and the complexities of many other models, the computational time complexity of a fuzzy logic regulatory model inference problem, particularly at higher order of interactions, quickly approaches those of computationally intractable problems. This undermines the benefits inherent in the simplicity and strength of the fuzzy logic-based molecular regulatory inference approach.

**Results:** For a sample inference problem – molecular regulation of vorinostat resistance in the HCT116 colon cancer cell lines, our modeled, designed and implemented “multistaged-hyperparallel” optimization approach significantly shortened the time to model inference from about 485.6 hours (20.2 days) to approximately 9.6 hours (0.4 days), compared to an optimized version of a previous implementation.

**Availability:** The multistaged-hyperparallel method is implemented as a plugin in the JFuzzyMachine tool, freely available at the GitHub repository locations https://github.com/paiyetan/jfuzzymachine and https://github.com/paiyetan/jfuzzymachine/releases/tag/v1.7.21. Source codes and binaries are freely available at the specified URLs.

## 1 Introduction

Fundamental to low- and high-throughput expression profiling is an enumeration of significant regulators of the biological processes under investigation. Among many known methods, the fuzzy logic approach appears most fascinating – simple, yet able to address or enunciate very complex relationships from expression profiles. Based on partial or imprecise classification of entities, the fuzzy logic describes an entity across multiple classifications. It ascribes a degree of membership for each possible class an entity may be classified [1,2]. Given a universe of objects, *U*, a subset or class of objects *A* can be described by applying a function (membership function, *f*) on a random selection of objects, *X* to derive a numeric value in the range [0,1]. An element, *x_i_*, in *X* can be said to belong to class *A* if derived value is greater than zero and the nearer the value of *f_A_*(*x*) to unity, the higher the ‘grade of membership’ of *x* in *A*. When *A* is a set in the ordinary sense of the term, its membership function can take only two values 0 and 1 [2]. In which case respective elements *x_i_*, in *X* are either not of or are of the class *A*.

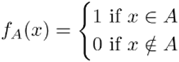

For a fuzzy set, different functions (membership functions) on *A*, *f_A_* can be considered – typically subjective and context dependent [3].

In its simplest description, fuzzy logic is the use of fuzzy sets in the representation and manipulation of vague information for the purpose of making decisions or taking actions [3]. It is a form of many-valued logic in which the truth values of variables may be any real number between 0 and 1 both inclusive, and employed to handle the concept of partial truth, where the truth value may range between completely true and completely false [4] contrasting Boolean logic (a two-valued logic), where the truth values of variables may only be the integer values 0 or 1 [5][6]. Many other specific examples do abound, including the Priest’s logic of paradox, the Bochvar’s internal three-valued logic, Belnap logic, Gödel logics, Product logic, Post logics, Rose logics, among others [7–9].

Though similar to probability in terms of range of value of between [0,1], fuzzy logic is not probability [3,10] because the forms of uncertainty addressed in both are different. The fuzzy logic jointly model uncertainty and vagueness unlike probability theory [11]. Bart Kosko argues that probability theory is a sub-theory of fuzzy logic [12]. Fuzzy logic extends classical logic to address uncertainty outside of classical logic and situations not amenable to probability theory.

With respect to regulatory processes, a fuzzy logic approach is thought to mitigate known challenges of modeling the biological system. These include inconsistencies and inaccuracies associated with high-throughput characterizations; challenges of dealing with noise and those of dealing with semi-quantitative data [13]. Like Boolean networks, fuzzy logic methods are simple and are fit to model imprecise and or highly complex networks [14,15]. But contrary to differential equation-based models, they are relatively less computationally expensive and less sensitive to imprecise measurements [14–16]. The fuzzy approach compensates for the inadequate dynamic resolution of a Boolean (or discrete) network, while simultaneously addressing the computational complexity of a continuous network [17]. With simple and easily scalable linguistic rules, a fuzzy logic-based inference system can make inference as would a trained expert looking into patterns of expressions among regulatory features.

Advantages with respect to using the fuzzy logic for expression dataset include; 1) An inherent account for noise in the data -- dealing with trends, not absolute values. 2) In contrast to other automated decision-making algorithms, such as neural networks or polynomial fits, algorithms in fuzzy logic are presented in the same language used in day-to-day conversations. Thus, fuzzy logic regulatory networks are more easily understood and can be extrapolated in predictable ways. And 3) fuzzy logic approaches are relatively computationally efficient and can be scaled to include an unlimited number of components [18].

A general fuzzy logic-based modeling and control system entails three major steps, 1) fuzzification, 2) rule evaluation and, 3) defuzzification [19].

1. Fuzzification: Considering expression as a linguistic variable and applying defined membership functions on observed expression data, the fuzzification step derives qualitative values – mapping non-fuzzy inputs to fuzzy linguistic terms [19]. A normalization technique may be applied to scale values to within a preferred range before fuzzification [17,19,20].
2. Rule evaluation: Driven by an inference engine, constructed rules in the form of “IF-THEN” are used to evaluate input variables and draw inference on the outputs alongside methods to aggregate results into definitive output. These include the maximum, bounded sum or normalized sum methods [19]. The fuzzy set operations (AND, OR, or NOT) are used to evaluate the fuzzy rules [1, 2]
3. Defuzzification: The defuzzification step produces a quantifiable expression result or value given the input sets, the fuzzy rules, and membership functions. Defuzzification technically interprets the membership degrees of the fuzzy sets into a specific decision or real value. The most common defuzzification approach is the center of gravity approach. This computes the center of gravity of the area under the membership function [21]. Where *X* is an ordinary non-void set, a mapping *A* from *X* into the unit interval [0,1] is the a fuzzy set on *X*, the value *A*(*x*) of *A* in *x* ∈ *X* is the degree of membership, the center of gravity defuzzification is given by [21]:

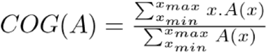

Other methods that are variants of the COG method include the basic defuzzification distributions (BADD) [22], mean of maxima (MeOM), indexed center of gravity (ICOG) among others [21].

Woolf and Wang (2000) presented one of the first applications of fuzzy logic to elucidate regulatory networks. Describing gene expression levels in linguistic terms of three possible states - low, medium, and high, they sort to find interacting gene triplets modeled as targets (T), activators (A), and repressors (R). Membership functions were employed to characterize expression levels as LOW, MEDIUM, and HIGH. With these, quantitative sets of rules were used to model regulatory networks. A sample predefined rule takes the form of “if A is LOW and R is HIGH, then T is LOW (Table 1). On each possible triplet, the expression linguistics were tested against the rules presented in the Table.

**Table 1:** Woolf and Wang’s rule table

|  | HIGH | MED | LOW |
| --- | --- | --- | --- |
| LOW | LOW | LOW | MED |
| MED | LOW | MED | HIGH |
| HIGH | MED | HIGH | HIGH |

In other words, their method entailed fuzzifying the expression data; creating and comparing gene triplets (activator-repressor-target) to generate a prediction value for the target (T) at points where the predicted values of A and R overlap; defuzzification to derive crisp values of target predictions; and triplet screening. Screening entailed comparing target predictions against observed expression values across biological experiments.

Almost immediately apparent is the computational complexity that is associated with Woolf and Wang’s approach - scaling in exponential time on the order *O*(*n*^3^) [18], where *n* is the number of interacting molecules. This quickly limits the number of interacting molecules that can be modeled to only three, i.e. two inputs and one output. Without improvement, the algorithm may only model simple regulation patterns and be unable to scale well to more complex models whose implementation time would be on the scale of years instead of hours [23]. Extending preliminary works of Reynolds [24], and by modifying the data preprocessing steps, Ressom and others did improve Woolf and Wang’s approach [25]. Although achieving an up to 50% improvement in computational time complexity over Woolf and Wang’s, we reason that Ressom and colleague’s preprocessing steps may tend to converge optimal models towards a local minimum. Also, the model inference time are still subjected to the problem of exponential growth in the number of rules to evaluate at higher orders of interacting molecules.

This problem of the exponential growth in the number of rules as inputs, compromising performance, associated with the intersection-rule configuration obtained in conventional fuzzy inference methodology of Woolf and Wang’s and others ([18,25,26] was addressed by Combs and Andrews [27]. They proposed an alternate rule configuration called the union-rule configuration (URC), together with a corresponding rule matrix called the union-rule matrix (URM) [28], to model the entire problem space without incurring any combinatorial penalty. Having first demonstrated the utility of the URC to qualitatively model the lac operon of E. coli [29,30], Sokhansaj et al extended the URC approach to model the yeast cell cycle from a time series expression data. In addition, their elucidated model was capable of qualitatively predicting data from another time series experiment [17].

To facilitate higher order model inferences in significantly faster computational time beyond Sokhansaj and others previously proposed methods, we performed a computational time complexity analysis of the fuzzy logic regulatory model inference system and, we developed and implemented a “multistaged hyperparallel” optimization approach. An alternate approach that achieves optimal model inferences in significantly faster computation time. In addition, it is generally agreed that fuzzy logic approaches to regulatory network inference can be scaled to include an unlimited number of components, but the associated time complexity benchmarked studies with available tools for future facilitation of comparisons is almost generally lacking. We anticipate this study would also provide both needed reference materials to benchmark future developments of the fuzzy logic-based regulatory model.

## 2 Methods

The methods in this study simply entailed; first, an analysis of Sokhansanj and colleagues optimized fuzzy logic method. Sokhansanj et al’s approach represent an improvement to the approach of Woolf and Wang, and that of Ressom et al’s. After an analysis of computational time complexity, we constructed a computation DAG (directed acyclic graph) associated with the algorithm. With the DAG, we estimated the algorithm’s ‘work’ and ‘span’ as described by Cormen et al. With values we obtained from the analysis of the DAG, we were able to describe efficiencies (theoretically, and validated empirically) using the Amdahl and Gustafson’s models. Furthermore, with the DAG, we were able to describe a multistaged hyperparallel approach to improve computation time inference.

### 2.1 The Fuzzy Logic Algorithm Analysis

As earlier mentioned, the impact of the computation algorithm employed can significantly affect the utility of the fuzzy logic approach to elucidate regulatory networks. The classical fuzzy logic triplet model of Woolf and Wang is reported to run on the order *O(n*^3^). Where *n* is the number of interacting molecules. This is a very conservative estimate as it accounts for only the number of fuzzy rule evaluations performed for a specific combination (activator-repressor-target) of triplet. It does not account for those of other combinations nor does it account for all other possible triplets. These can have a combinatorial explosion-like growth function that quickly become significant in comparison to that observed with the rules evaluated with increasing *n*. Employing the union-rule configuration (URC), Sokhansanj et al were able to reduce the complexity of Woolf and Wang’s solution from *O*(*m^NN^*) to *O*(*m^N^*). Where *N* is the number of (input) genes regulating an output gene and *m* is the number of possible rules describing the effect of each single input gene on an output gene. The number of possible rules for each gene-gene interaction (*m*) is given by *n^n^*, where n is the number of fuzzy sets that describe the state of a variable [17]. Similarly, this is a very conservative estimate. It accounts for only the number of fuzzy rule evaluations performed for a specific combination of a particular set of inputs (regulators) and output genes. It does not account for those of other combinations of input genes nor does it account for all other possible combinations of inputs (regulators) and output genes which may similarly exhibit a combinatorial explosion-like growth function.

#### 2.1.1 Theoretical Analyses of computational time complexity

To analyze the computational time complexity of the Sokhansanj approach, a pseudocode is presented here

#### 2.1.2 The Exhaustive search

~~~
1. Read-in the configuration file
2. Initialize object
3. Initialize table of expression
4. Initialize fuzzy Matrix (fuzzified values of expression values)
5. // to enable a constant-time access to fuzzy sets of expression values
   // do exhaustive search:
6. get the output nodes (output genes), *ON*_1_, *ON*_2_, *ON*_3_···*ON_N_*
7. // these may be all the genes in expression matrix or a pre-specified number
8.        for each of the output gene node:
9.         get other genes (potential inputs to the current output node)
10.        get the ‘number of inputs’ nodes to consider
11.          // may consider a maximum number of input nodes *IN_Max_*
12.          // defaults to a specific user specified number of inputs
13.          // do deeper search:
14.          get the desired e-value cut-off (e-cutoff)
15.          get combinations (permutations) of input nodes; *CIN_1_···CIN_p_*
16.          get output gene expression values
17.          get mean expression value of output gene
18.          get the sum of squared deviations (dss) of output gene expression values
19.          get combinations of inputs
20.          for each combination (of inputs):
21.             get all possible combinations of fuzzy rule to evaluate
//nested for loops
22.             for each possible combinations of fuzzy rule
23.                 instantiate a string array for the input genes
24.                 // compute residuals
25.                 for each expression value of the output gene across all samples, time series or perturbations
26.                     get input genes
27.                     get the fuzzy set values for respective input genes
28.                     perform a union rule configuration (URC) evaluation
29.                     defuzzify aggregate fuzzy set
30.                     compute residual squared sum (rss)
31.                     // sum squared residual
32.                     compute fit (error) = 1
33.                     if computed fit is greater than or equals e-cutoff
34.                         populate fuzzy ‘rule’ arrays with valid rule instances…
35.                         instantiate a result object
36.                         // may add result object into a collection of result objects
37.                         print acceptable result
38.                     end if
39.                  end for
40.               end for
41.            end for
42.          end for
~~~

From a set of output nodes (gene features to be included in the derived regulatory network), the algorithm independently and exhaustively searches for models (combinations of inputs to output), across samples, that meet prespecified fit cut-off (lines 6 - 42). From a calculation of operations in the outlined pseudocode, time complexity is approximately:

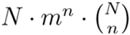

Where

*N* is the number of output genes being considered.
*m* is the number of possible rules for each gene-gene interaction, this is the square of the number of fuzzy sets that describe a variable. For a three-fuzzy-sets (LOW, MEDIUM, and HIGH) model, this would be 3^3^, which is 27. And,
*n* is the number of input nodes being considered.

### 2.2 Analyzing ‘work’ and ‘span’

The fuzzy logic algorithm consists of solving many smaller, independent comparisons; lending itself to parallel computing and anticipated to potentially scale linearly with the number of available compute processors [18]. In recent times, more readily available workstations with much faster clock-speeds and multicore/multithreaded abilities, including ready access to high performance computing environments present opportunities to investigate larger regulatory inference problem sizes such as those of higher degrees of interactions at individual nodes of a regulatory network with the fuzzy logic model. For a theoretical analysis of a potential many-processors solution, we described a computation directed acyclic graph (DAG) from the previously outlined algorithm (pseudocode) and we defined two metrics - “work” and “span”, as specified by Cormen et al [36]. Work is defined as the total time to execute the entire computation on one processor. For our computation DAG in which each edge is assumed to take a unit time, work is equivalent to the total number of vertices. Span is the longest time to execute the strands along any path in the DAG. A strand is a chain of instructions containing no parallel control, in the DAG. For our DAG, the span equals the number of vertices on a longest or critical path in the DAG (colored path in Supplementary Figure 1). For our example computation DAG, the total number of vertices would be given as:

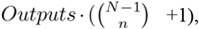

Where,

*Outputs* is the number of output genes being considered,
*N* is the number of genes in the network, and
*n* is the number of input genes being considered.

For a single output and two input genes from a fifty gene set, our computation DAG would have approximately 1177 vertices (work of 1177 time units, see results and discussion section) and a span of 2 vertices (2-time units).

### 2.3 Measuring performance (theoretical efficiency)

To theoretically estimate efficiency, we employed both Amdahl and Gustafson models. Gene Amdahl on what eventually became known as Amdahl’s law made a submission for a single processor approach to large scale computing, arguing that for most applications, there exist a sequential potion that cannot be parallelized. He argued that, with an increasing number of processors, this sequential portion may constitute up to 50% - 80% of the total execution time and thus have a diminishing effect. Also referred to as the fixed-size speed up model, Amdahl’s law states that if a portion of a computation, *f*, can be improved by a factor *m*, and there exists another portion that cannot be improved, then the portion that cannot be improved will quickly dominate the performance, and further improvement of the improvable portion will have little effect [31–34]. This is given by:

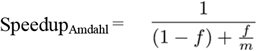

Where *f* is a parallelizable portion, and *m* the number of processors; as m→∞,

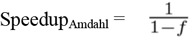

However, together with colleagues at the Sandia National Laboratories working on a 1024 processor system, Gustafson and colleagues demonstrated that the assumptions of Amdahl’s argument were inappropriate to describe observed results with massive parallelism [31–34]. Identifying the shortfall in an implicit assumption in Amdahl’s law - that the number of processors is independent of size of the problem, Gustafson proposed that it would be more realistic to assume run time, not problem size as constant. This was subsequently referred to as the fixed-time speedup model [32] (SpeedupFT), and is given as:

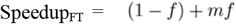

Known as the Gustafson’s law, it is a linear function of *m* if the workload is scaled to maintain a fixed execution time.

From our analysis, the fractional part of the fuzzy algorithm that cannot be made parallel relative to the entire computation is the ratio of estimates of ‘span’ to ‘work’ described above.

### 2.4 The “Multistaged Hyperparallel” Optimization

The approach simply divides the inference problem to its many computable units, grouped at two stages in its computation directed acyclic graph (DAG) (see section below, and Supplementary Figures 1 and 2).

#### 2.4.1 Multistaging

With respect to modelling parallel executions, Cormen et al suggested that it helps to think of a computation as a directed acyclic graph *G* = (*V, E*), called a computation DAG (Supplementary Figure 1). Conceptually, the vertices in *V* are instructions and data objects, and the edges in *E* represent dependencies between instructions and data objects, where (*u, v*) *∈ E* means that the set of instruction *u* must execute before instruction *v* [36]. Logically, if a vertex v has a direct path from another vertex *u*, both processes, *u* and *v* are described as (logically) in series but logically in parallel if not [36].

Practically, a closer examination of outlined fuzzy logic algorithm pseudocode shows multiple lines of dependent instructions and blocks of potentially parallel operations (Supplementary Figures 1 and 2). It is compelling that computable units can be grouped at two stages (Supplementary Figure 2), at a very least. Also compelling is that Staged I computable units (Figure 2) can simultaneously be presented to a job scheduler like SLURM [40,41] or similar schedule manager within a high-performance computing (HPC) environment, to maximize a breaking down of the inference problem and to achieve a distribution across many more processor cores not only in parallel but in a parallel-parallel (hyperparallel) manner within a shorter time frame. This may represent a re-formulation of the inference problem as one which conforms to Gustafson’s rather than Amdahl’s model – increasing core requirement as a function of the problem size (in this case, the number of output nodes to consider), in order to aim for a fixed execution time as seen with a consideration of a single output node.

The SLURM scheduler is a de facto manager on many HPC environments which facilitates dynamic multithreading (parallel processing) and allowing computation to specify parallelism without worrying about communication protocols between environment nodes, load balancing, and other peculiarities of static threads [40, 41].

## 3 Results and Discussions

Given the following number of inputs, and fifty output nodes, analytical estimates of computational time complexity are estimated in Table 2.

**Table 2:**
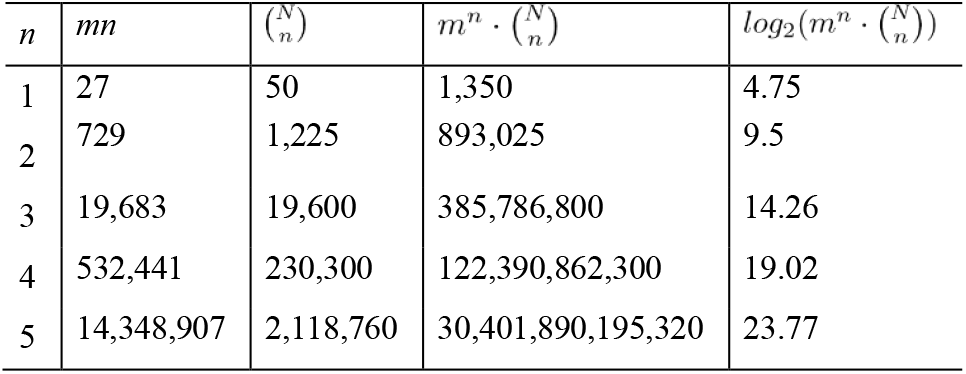
Theoretical time complexity estimates. Analytical estimates of computational time complexity in milliseconds. Note that the total number of outputs, *N*, being constant, was omitted in estimating the big *O*.

| $n$ | $mn$ | $\binom{N}{n}$ | $m^n \cdot \binom{N}{n}$ | $\log_2(m^n \cdot \binom{N}{n})$ |
| --- | --- | --- | --- | --- |
| 1 | 27 | 50 | 1,350 | 4.75 |
| 2 | 729 | 1,225 | 893,025 | 9.5 |
| 3 | 19,683 | 19,600 | 385,786,800 | 14.26 |
| 4 | 532,441 | 230,300 | 122,390,862,300 | 19.02 |
| 5 | 14,348,907 | 2,118,760 | 30,401,890,195,320 | 23.77 |

Supplementary Figure 3 shows a plot of the logarithm of the analytical estimate of time complexity (calculated cost) as a function of inputs to the algorithm. The derived log estimates were fitted using a simple linear regression model to obtain a slope, estimated to be 8.2146. This implies that for every additional input considered, the computational time complexity grows by approximately eight folds, if every other variable or factor remains constant.

### 3.1 Empirical analyses of computational time complexity

To investigate how well our estimates, capture real world situations, we ran our implementation of the algorithm with up to four inputs. Table 3 shows the execution time, considering only one output node. Aside from the differing number of input nodes considered, all other factors remained the same. The computation experiment was performed on a compute node (x86_64 Genuine Intel Xeon Gold 6150 CPU@2.70GHz) with 32 cores and 18GB of available runtime memory on the Frederick National Laboratory High Performance Computing Cluster Environment.

**Table 3:**
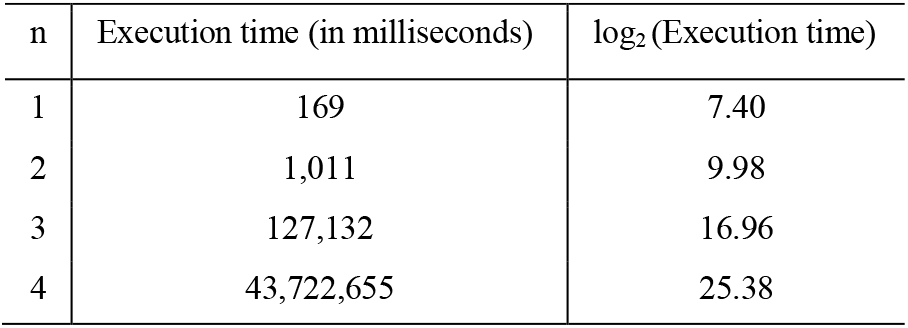
Empirical execution time

We also fitted the observed log value of execution time using a simple linear regression model to obtain a growth rate, i.e. slope (Supplementary Figure 4). Though empirical growth rate of the fitted curve appears to be less than that estimated from a complexity analyses of the algorithm (Supplementary Figure 3), the nature of the empirical curve growth rate beyond two inputs appears almost parallel to that of the analytical estimates. We previously mentioned that a *m^n^* computation time complexity specification underestimates the real-world nature of the fuzzy logic algorithm and here our estimation show that such is the case – Supplementary Figure 5 shows the slope of *m^n^* curve diverging away from the fitted curve of empirical observation, underestimating computational time as the number of inputs increases. Supplementary Figure 6 shows a plot of the theoretical or expected speedup that can be achieved at varying number of available computing cores in a high-performance computing (HPC) environment, not considering node, scheduling, I/O, memory or other possible computational overheads. The figure shows theoretical speedup for both Amdahl and Gustafson’s models. Speedup estimate using Gustafson’s model does appear to increase linearly with available computing cores. However, it appears to approach an asymptotic maximum with Amdahl’s model.

Cormen et al described “parallelism” of a multithreaded (and by extension a parallel) computation as the average amount of work that can be performed in parallel for each step along a critical path. Its estimation is described as the maximum (upper bound) speedup that can be achieved on any number of processors [36]. Given that work estimates from our computation DAG (Figure 1) is T1 = 1177 and span (irrespective of the number of available processors) is T∞ = 2. Cormen et al defined parallelism as T1/T∞, which approximates to 589. From Supplementary Figure 6, this very closely mirror a possible speedup upper bound estimate from Amdahl’s model.

**Figure 1.**
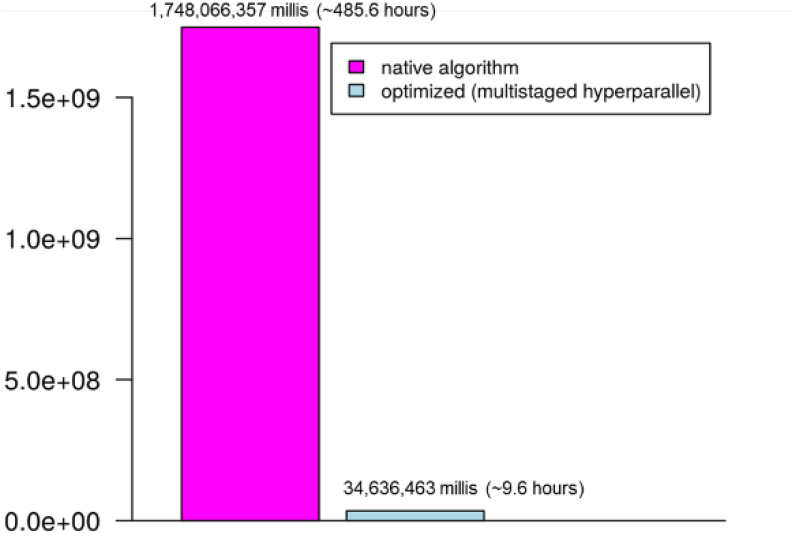
Comparison of execution time (in milliseconds) between native algorithm and optimized (multistaged hyperparallel) algorithm. For a sampled inference problem, the “multistaged hyperparallel” optimization approach is demonstrated to significantly shorten time to model inference from about 485.6 hours (20 days) to approximately 9.6 hours (0.4 days).

Measuring performance (empirical efficiency) of our optimized fuzzy logic algorithm and to evaluate how well our theoretical evaluation mirrors a real-world situation. We ran our algorithm with the same parameters (a single output gene, two input regulatory genes and a fifty genes set) on a multicore machine (64bit x86_64, Genuine Intel CPU@1.80GHz) and observed the execution time using 1 to11 computing cores; where T1 is the algorithm’s execution time with just one core and TP is the execution time on a specified number of processors *p*, Table 4 shows observed execution time in milliseconds while Supplementary Figure 7 is a representative bar plot of the execution times. Supplementary Figure 8 shows a plot of speed up versus available computation core (processor). Demonstrating that computation time complexity does scale linearly, the fitted line of the plot shows an almost linear growth curve. The largest change in speedup gradient appears to be between one and two cores.

**Table 4:**
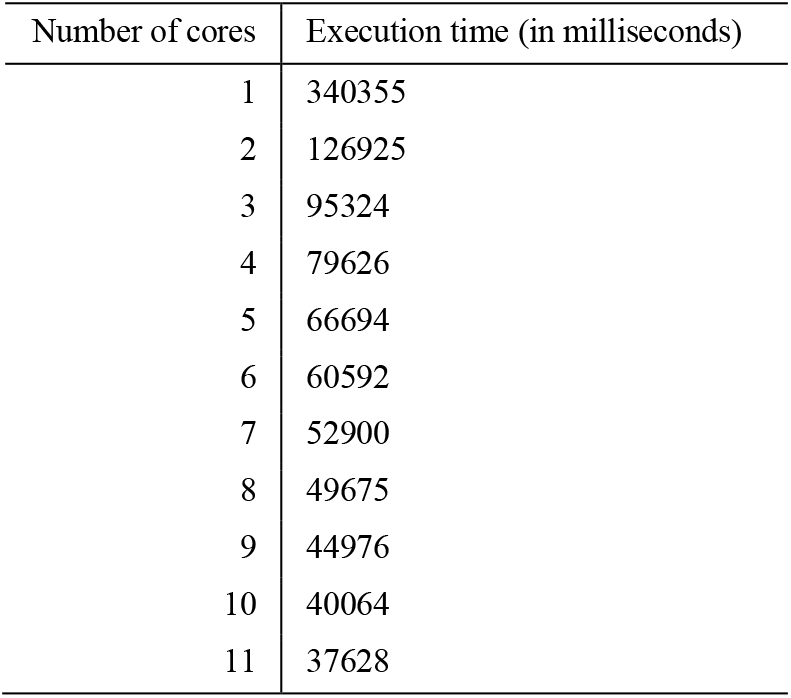
Empirical execution time

Supplementary Figure 9 overlays Amdahl’s and Gustafson’s model speedup estimates at the respective number of cores. Empirically observed speedup at one to four processing cores appear higher that both predictions of Amdahl and Gustafson speedup models. However, the generally slower rate of change of the curve quickly brings the speedup gain per increase in core to below Amdahl and Gustafson’s. At a lower number of processing cores, both Amdahl’s and Gustafson’s estimates appear to trend together, however these begin to diverge at about 8 or 9 processing cores.

To show the increase in efficiency obtained with the multistaged hyperparallel approach, we applied it to a sample Fuzzy logic based regulatory network inference problem – one with 50 separate output genes, 4 input regulatory genes from a fifty-genes set. From Figure 1, the “multistaged hyperparallel” optimization approach is demonstrated to significantly shorten time to model inference from about 485.6 hours (20.2 days) to approximately 9.6 hours (0.4 days).

Representing an almost 50 fold increase in speedup, which almost correspond to the number of output genes being considered and also corresponds to the number of Stage I grouped computation units (see Figure 2, and section on multistaging), the multistage hyperparallel approach tend to keep execution time constant for every increase in output genes considered to gain a corresponding fold speedup, provided all other parameters remain the same. It may well be assumed that the multistaged hyperparallel approach tend to reformulate the fuzzy logic computation and inference problem from that which obeys the Amdahl’s law to one which approximates the Gustafson’s model.

## 4 Conclusions

The fuzzy logic regulatory network inference method is a simple yet powerful approach to elucidating interacting molecule in regulatory networks whose efficiency becomes undermined by high computational complexity at higher order interacting molecule inference problems. The multistaged hyperparallel approach presented and this benchmark study demonstrates that though the fuzzy inference system is amenable and readily scales with additional compute cores, the speedup gained per unit increase in compute core, within a high-performance computing environment diminishes and more likely to approach an asymptotic maximum, tending to more closely mimic Amdahl’s model than the Gustafson’s model. The multistaged hyperparallel optimization approach presented however, significantly improves computation time, by reformulating, in practical terms, the inference problem from what follows the Amdahl’s to that which approximate Gustafson’s model.

## 5 Availability

The source code and binaries of the optimized fuzzy logic, with the union rule configuration, regulatory network inference engine, including enabling the multistaged-hyperparallel configuration are implemented in the JFuzzyMachine tool available at the bitbucket git repository locations https://bitbucket.org/paiyetan/jfuzzymachine/src/master/ and https://bit-bucket.org/paiyetan/jfuzzymachine/downloads/. Source codes and binaries are available free for academic and non-commercial use.

## Acknowledgements

A special thanks to Dr. Iosif Vaisman, Dr. Dmitri Klimov and Dr. Saleet Jafri, at the George Mason University School of Systems Biology and Dr. Andrew Quong (formally at the Frederick National Laboratory for Cancer Research, and now at the Fluidigm Corporation), for mentoring the larger body of the work for which a component part is presented here. Also, the contribution of the high-performance computing resources at the Frederick National Laboratory for Cancer Research (FNLCR) and the Extreme Science and Engineering Discovery Environment (XSEDE) supported by the National Science Foundation at the Texas Advanced Computing Center (TACC) at The University of Texas at Austin - http://www.tacc.utexas.edu, are greatly acknowledged.

## Funding

The Frederick National Laboratory for Cancer Research is fully funded by the National Cancer Institute. This project has been funded in part with Federal funds from the National Cancer Institute, National Institutes of Health, under contract number HHSN261200800001E. The content presented here is solely the responsibility of the author and does not reflect the official views of the National Institute of Health, the National Cancer Institute nor the Frederick National Laboratory for Cancer Research.

## Conflict of Interest

none declared.

## Notes

### Competing Interest Statement

The authors have declared no competing interest.

